# *Ex Vivo* and *In Vivo* CD46 Receptor Utilization by Species D Human Adenovirus Serotype 26 (HAdV26)

**DOI:** 10.1101/2021.09.28.462271

**Authors:** Jack R. Hemsath, A. Manuel Liaci, Jeffrey D. Rubin, Brian J. Parrett, Shao-Chia Lu, Tien V. Nguyen, Mallory A. Turner, Christopher Y. Chen, Karolina Cupelli, Vijay S. Reddy, Thilo Stehle, M. Kathryn Liszewski, John P. Atkinson, Michael A. Barry

**Affiliations:** Graduate Research Employee Program (GREP), Mayo Clinic, Rochester, MN 55905, USA; Virology and Gene Therapy Graduate Program, Mayo Clinic, Rochester, MN 55905, USA; Division of Infectious Diseases, Department of Medicine, Department of Immunology, Department of Molecular Medicine, Mayo Clinic, Rochester, MN 55905, USA; Interfaculty Institute of Biochemistry, University of Tuebingen, D-72076 Tuebingen, Germany; Department of Integrative Structural and Molecular Biology, The Scripps Research Institute, La Jolla, CA, 92037, USA; Division of Rheumatology, Department of Medicine, Washington University School of Medicine, St. Louis, MO 63110; Structural Biochemistry, Bijvoet Centre for Biomolecular Research, Utrecht University, Universiteitsweg 99, 3584 CG, Utrecht, The Netherlands

**Author notes:** **Correspondence to:** Michael A. Barry, PhD., Mayo Clinic, 200 First Street SW, Rochester, MN, USA.

## Abstract

Human adenovirus serotype 26 (Ad26) is used as a gene-based vaccine against SARS-CoV-2 and HIV-1. Yet, its primary receptor portfolio remains controversial, potentially including sialic acid, CAR, integrins, and CD46. We and others have shown that Ad26 can use CD46, but these observations were questioned by the inability to co-crystallize Ad26 fiber with CD46. Recent work demonstrated that Ad26 binds CD46 with its hexon protein rather than its fiber. We examined the functional consequences of Ad26 for infection *in vitro* and *in vivo*. Ectopic expression of human CD46 on Chinese hamster ovary cells increased Ad26 infection significantly. Deletion of the complement control protein domains CCP1 or CCP2 or the serine-threonine-proline (STP) region of CD46 reduced infection. Comparing wild type and sialic acid-deficient CHO cells, we show that the usage of CD46 is independent of its sialylation status. Ad26 transduction was increased in CD46 transgenic mice after intramuscular (IM) injection, but not after intranasal (IN) administration. Ad26 transduction was 10-fold lower than Ad5 after intratumoral (IT) injection of CD46-expressing tumors. Ad26 transduction of liver was 1000-fold lower than Ad5 after intravenous (IV) injection. These data demonstrate the use of CD46 by Ad26 under certain situations, but also show that the receptor has little consequence by other routes of administration. Finally, IV injection of high doses of Ad26 into CD46 mice induced release of liver enzymes in the bloodstream and reduced white blood cell counts, but did not induce thrombocytopenia. This suggests that Ad26 virions do not induce direct clotting side effects seen during COVID-19 vaccination with this serotype of adenovirus.

**IMPORTANCE:** Human species D Ad26 is being pursued as a low seroprevalence vector for oncolytic virotherapy and gene-based vaccination against HIV-1 and SARS-CoV-2. However, there is debate in the literature about its tropism and receptor utilization, which directly influence its efficiency for certain applications. This work was aimed at determining which receptor(s) this virus uses for infection, and its role in virus biology, vaccine efficacy, and importantly, in vaccine safety.

## INTRODUCTION

Adenoviruses (Ads) are a genetically diverse family that cause a range of ocular, respiratory, digestive, and systemic infections (1). Human mastadenoviruses (HAdVs) are grouped into species A through G with whole genome sequence diversity as high as 40% from one end of the virome to the other (2). Despite this drastic genetic diversity, most Ad biology is extrapolated from the human species C adenovirus Ad5, possibly due to historical factors and the unavailability of other Ad serotypes commercially.

Human species D Ad26 (HAd-D26) is currently being pursued as a gene-based vaccine and as an oncolytic virus for use in humans (3-11). Ad26 has most recently been used as replication-defective adenovirus (RD-Ad) vaccines against SARS-CoV-2 (12, 13). It has been demonstrated that intramuscular (IM) vaccination with high doses (10^11^ viral particles (vp) of Ad26 expressing different spike proteins could generate protection against respiratory challenge with SARS-CoV-2 challenge in macaques. While Ad26 has utility as an HIV-1 and COVID-19 vaccine, there is debate in the literature about this virus’ receptor utilization and tropism.

Most Ad tropism is dictated by the interactions of their fiber and penton base proteins. Ads use their fiber proteins to bind several receptors including the coxsackie and adenovirus receptor (CAR), sialic acid, and CD46 (reviewed in (14, 15)). The archetype Ad5 virus and its species C family members Ad1, 2, 6, and 57 bind CAR. In contrast, many species B Ads such as Ad21 and Ad35 bind CD46 (16-19) Most (but not all) adenoviruses also bind integrins as primary or secondary receptors using RGD motifs in their penton base (20-22). Furthermore, many species D and G Ads recognize sialic acid-decorated proteins or glycolipids. In most cases, fiber mediates initial docking to cells, and lower affinity integrin interactions then mediate secondary binding and cell entry by endocytosis (23). Ads can exclusively use their penton-integrin interactions if the target cell lacks fiber receptors, or when fiber affinity for receptors is low (23).

Species D Ads are currently the largest group of adenoviruses in the human virome. For most, it is unclear what primary disease they may cause. A subset of species D Ads including Ad37 cause severe keratoconjunctivitis in humans (24, 25). Species D viruses access a diverse portfolio of receptors including CAR, sialic acid, and CD46, but the specific arsenal varies between serotypes and is often difficult to assess (reviewed in (14, 15). Species D Ads have short, stiff fibers with only 8 repeats (**Fig. 1**and (26)) making CAR interactions less efficient than in other species (14). The Ad37 fiber binds to sialic acid, with high selectivity for the GD1a glycan motif (24, 25). While Ad37 binds sialic acid, the affinity of this interaction is only 19 µM which is approximately 1,000-fold lower than the affinity of the Ad5 fiber for CAR and the Ad11 fiber for CD46 (17, 27).

**Figure 1.**
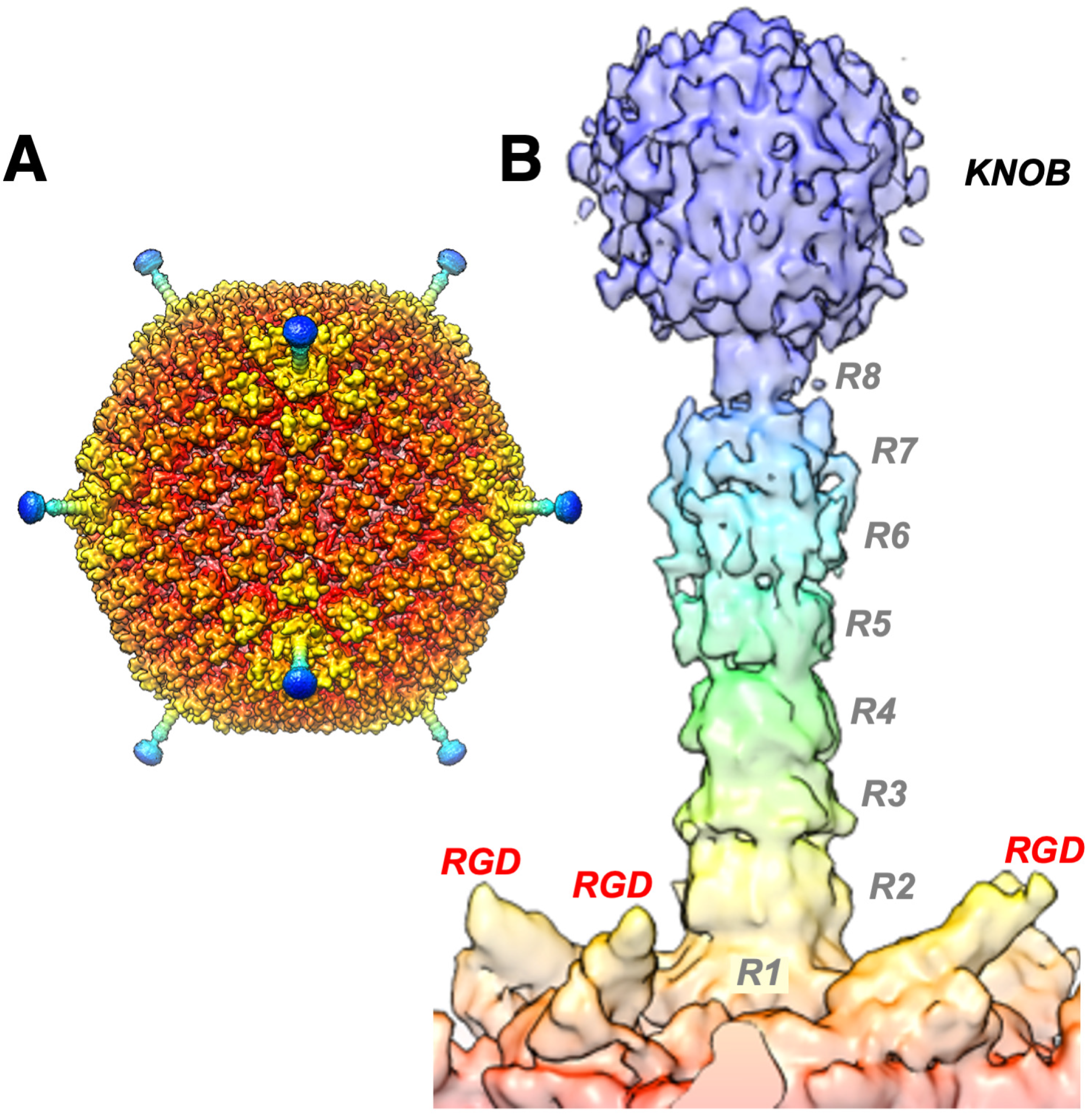
Cryo-Electron Microscopic Structures of Ad26. **A)** Full virion structure **B)** Fiber and penton base. R indicated fiber shaft repeats. RGD indicates arginine-glycine-aspartic acid integrin binding motifs in the penton base. Knob indicates the receptor binding portion of the Ad26 fiber trimer.

CD46 or membrane cofactor protein is a widely expressed membrane-bound complement control protein ((28, 29) and reviewed in (30, 31). CD46 has an extracellular ectodomain containing four complement control protein domains (CCP1 to CCP4) that are also known as short consensus regions (SCR1 to SCR4) on its N-terminus (**Fig. 2** and (30)). The four CCPs are attached to an alternatively spliced variable-length serine-threonine-proline (STP) rich region, a transmembrane domain and two alternate cytoplasmic tails (28). The four CCPs form an extended “hockey stick” structure that allows different interactions with complement and pathogen proteins (**Fig. 2** and (32)). This structure also allows CD46 to display N- and O-linked carbohydrates on its CCP and STP domains, respectively, that can be capped with sialic acids, potentially mediating additional interactions with sialic-acid binding Ads. (**Fig. 2** and (33)). Most crystallographic data on Ad knob-CD46 interactions has been obtained using the N-terminal CCP1 and CCP2 domains of CD46 (known as CD46-D2) (19).

**Figure 2.**
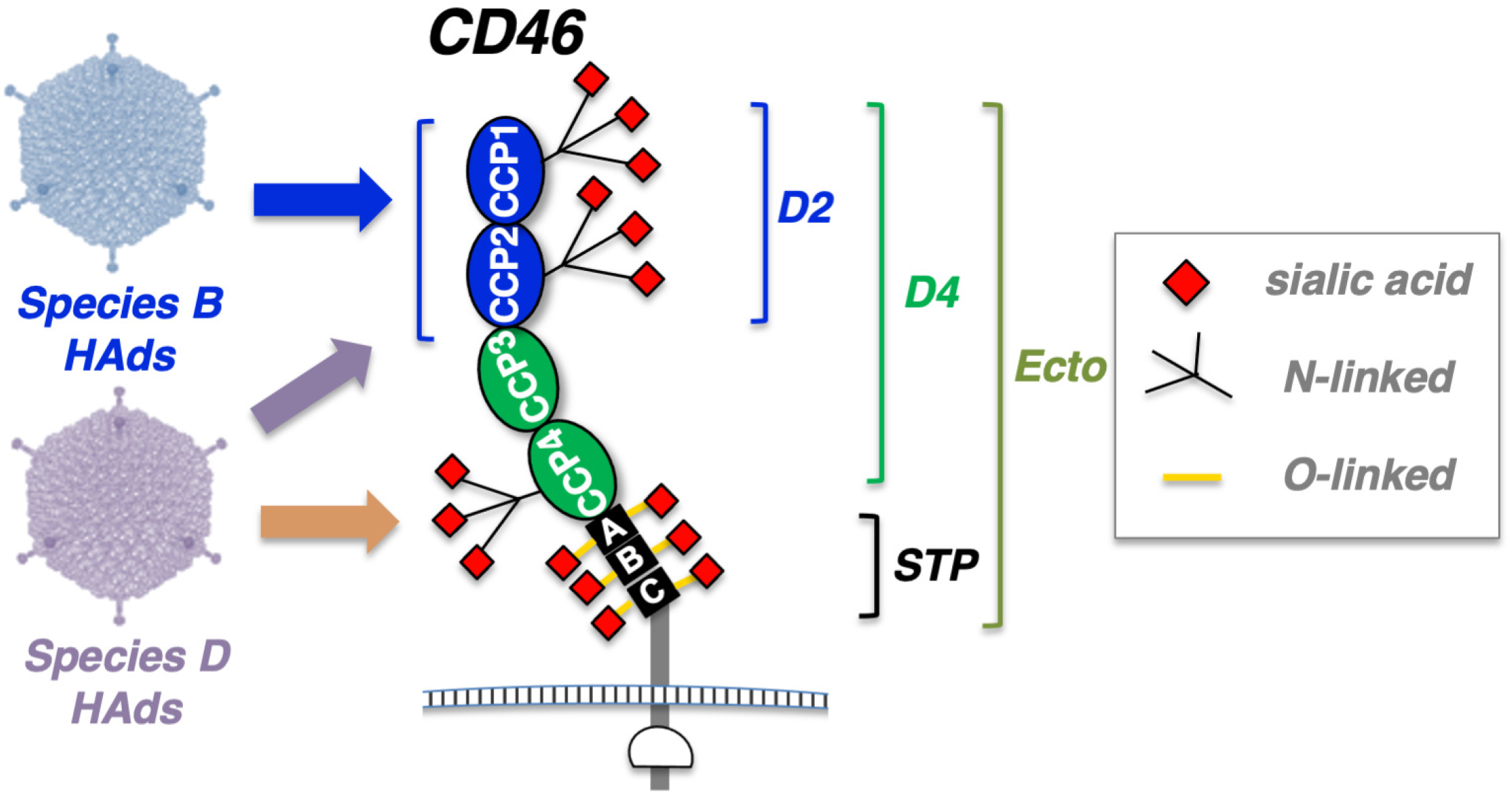
Schematic of Potential Adenovirus Binding Motifs on CD46. CCP, STP, D2, D4, Ecto, domains are described in the text. Species B Ads are known to bind the CCP1 and CCP2 domains on CD46.

The CD46 transcript is alternatively spliced to produce four predominant protein isoforms on most cells (28, 34, 35). All forms contain the four CCPs but vary in the number of STP domains. Most tissues express one to two STP domains (BC or C) and one of two different cytoplasmic tails (CYT-1 or CYT-2). The four common isoforms are named BC1, BC2, C1, and C2. All four isoforms co-exist on most human tissues, but in differing proportions. Many tissues express the BC and C isoforms (28, 34-36). BC2 isoforms predominate in the kidney and salivary glands (35). C2, in turn, is expressed in the brain (35), and a hypoglycosylated form is observed in spermatozoa (37, 38). A fifth CD46 variant including all three STP domains, named ABC, is rarely observed (28) and expressed at low levels in lung, ovary, placenta, and testes. Additionally, few tissues (brain and bladder) can express CD46 lacking all STP domains (36). While CD46 is broadly expressed on human cells, this is not the case in mice. Mice do not express CD46 in most tissues, but instead express it only in the testes and in the retina (37, 39-42).

Different pathogens use different domains of CD46 for cell binding (reviewed in (30)(15)). Measles virus uses CCP1 and CCP2, human herpes virus 6 uses CCP2 and CCP3, *Streptococcus pyogenes* uses CCP3 and CCP4 and *Neisseria gonorrhoeae* and *Neisseria meningitidis* use CCP3 and the STP domain. Species B adenoviruses bind the CCP1-CCP2 domain of CD46 (CD46-D2) using the knob domain of their fiber protein (**Fig. 2** and (14, 19)).

Initial *in vitro* data on artificial CAR and CD46-modified B16F10 cells suggested that Ad26 did not use CAR, but instead used CD46 as a receptor for infection (3). While Ad26 infection was increased on cells that artificially expressed CD46, the level of infection by Ad26 on these cells was less than half as efficient as infection by CD46-utilizing species B Ads Ad11 and Ad35 (3). Subsequent work by our laboratory on primary human patient B cell cancer cells inferred that Ad26 used a mixture of CD46 and integrin binding to infect these cells (7). CD46 blocking antibodies reduced Ad26 infection in these B cells by only 50%. This blockade could be increased to 100% by also blocking binding to integrins with cyclic RGD peptide (7). While species D Ads can use sialic acid as a receptor, Ad26 infection was unaffected by removing or blocking sialic acid on these cells. Ad26 is markedly more effective at infecting CD46 transgenic mice than normal mice after intramuscular injections (43).

These data suggested that Ad26 uses CD46 and integrins as receptors, albeit not as efficiently as other Ads. A more recent study reports that Ad26 does not use CD46 at all when infecting epithelial cells, but instead uses αvβ3 integrin as its primary receptor (44). These well-controlled studies showed that knock-down of CD46 with siRNA did not inhibit Ad26 infection. They also showed that CHO cells expressing the BC isoform of CD46 are not infected at higher efficiencies than CHO cells (44). This observation using CD46-BC expressing cells conflicts directly with earlier data testing Ad26 on B16F10-CD46 cells (3). It is unclear from this earlier report which isoform of CD46 is expressed on the B16F10 cells, so it is uncertain if this is a source of the discrepancy.

Another recent study from Baker *et al*. crystalized the Ad26 knob and performed structural and biological analyses of its interactions with CAR, CD46, and desmoglein-1 (45). This work showed that the Ad26 knob had 20 µM Kd for CAR and 50 µM Kd for CD46-BC1 ectodomain by surface plasmon resonance (SPR) (45). Co-crystals could only be produced with CAR, but not with CD46 or desmoglein-1. Subsequent work by the same group showed that removing sialic acid from cells inhibits Ad26 infection and that its knob could be co-crystalized with sialic acid (46). From this, it was concluded that CD46 was not the receptor for Ad26.

The conundrum of observations of Ad26 using CD46 versus the failure of the Ad26 knob to bind or co-crystalize with CD46 was resolved by recent seminal work (47). This work demonstrated that Ad26 does indeed bind CD46, not with its fiber, but rather with its hexon protein.

Given the utility of Ad26 for vaccination and cancer therapy, but the uncertainty in its receptor utilization, we have further explored the ability of Ad26 to use CD46 as a receptor *in vitro* and importantly *in vivo*. This work also tested the possibility that the CD46 protein itself might not be the receptor for Ad26, but that instead CD46 might simply be serving as a scaffold to display sialic acid for the virus to bind (**Fig. 1**). In this case, we hypothesized that Ad26 might not only bind CCPs on CD46 but might also or instead bind to sialic acids on the CCPs or on the heavily *O-*glycosylated STP domain of the protein.

This work compares the *in vivo* activity of Ad26 with benchmark Ad5 virus in mice with and without CD46. Finally, given recent concerns with thrombotic thrombocytopenia in people vaccinated with Ad26 and ChAdOx1 (48, 49), this study also examined if intravenous delivery of high doses of Ad26 into CD46 transgenic mice can provoke changes in blood counts, platelets, and blood chemistry.

## MATERIALS AND METHODS

### Cell Lines

Cells were maintained in HyClone Dulbecco’s High Glucose Modified Eagles Medium (Thermo Scientific, Waltham, MD) supplemented with 10% fetal bovine serum, 100 U/mL penicillin and 100 µg/mL streptomycin. 293 and 293-IX-E4 cells were purchased from Microbix (Mississauga, Ontario, Canada). Chinese hamster ovary (CHO) and Lec2 cells were purchased from American Type Culture Collection (ATCC, Manassas, VA). CHO-CD46-C1, CHO-CD46-BC-1, CHO-CD46-BC-1-ΔCCP1, CHO-CD46-BC-1-ΔCCP2, CHO-CD46-ΔSTP, CHO-CD46-C2, and CHO-CD46-BC-2 cells were generously provided by Dr. Kah-Whye Peng (Mayo Clinic) or by Dr. Kathy Liszewski and Dr. John Atkinson. Human CD46-BC1 cDNA (accession BC030594) was purchased from Transomic Technologies (Huntsville, AL, USA). pCDNA-CAR and pCDNA-JAM1 plasmids were generously supplied by Dr. Terrence Dermody (University of Pittsburgh). CHO and Lec2 cells were transfected with plasmids expressing human CD46-BC1 (accession BC030594), CAR, or Junctional Adhesion Molecule 1 (JAM1) using Polyfect (Qiagen, CA, USA) and selected in 1000 µg/ml G418 for stable transfectants. The CD46 ecto domain (Ecto) was PCR’d off the human CD46-BC1 BC030594 cDNA using CD46 5 prime SCR (CCAGATCTCTGTGAGGAGCCACCAACATTTGAAG) and CD46 3 prime STP (CTCTCTGGGCCCTCTAGATGAGACTGGAGGCTTGTAAGTAGG). The CD46 BC STP domain was PCR’d with CD46 5 prime STP (CCAGATCTCAAAGTGTCGACTTCTTCCACTACAAAATC) and CD46 3 prime STP. These were cut with BglII and ApaI and cloned into pHook2-PSTCD (50) in frame and between the alpha1-antitrypsin secretory leader and the PDGFR transmembrane domain from pHook2 (Invitrogen) that allows cell surface display of fusion proteins. CHO and Lec2 stable cell lines were selected with G418 as above. Stable cells were stained with MEM-258 CD46 antibody and verified on a flow cytometer for CD46 expression before use (Supplemental Figure S1). B16 mouse melanoma cells were purchased from ATCC. These cells were first modified with pCDNA3-CAR expressing human CAR by transient transfection and selection with G418. B16 cells were also transduced with a lentivirus expressing human CD46 followed by selection in puromycin. CAR and CD46 expression in the resulting B16-CAR-CD46 cells was verified by flow cytometry.

### Adenoviruses

Replication-defective Ad5 (RD-Ad5) and replication-competent Ad5 (RD-Ad5) expressing the green fluorescent protein-luciferase (GFP-Luc) fusion protein were produced as described in (51). RD-Ad6, RD-Ad26 and RC-Ad26 expressing GFP-Luc were generated as in (52). RD-Ad5, RD-Ad6 and RC-Ad26 were grown in 293 cells. The Ad5 E1 in 293 cells supports E1 deleted species C viruses, but not other Ads species viruses (53). Therefore, E1-deleted RD-Ad26 was grown in 293-IX-E4 cells that also provide Ad5 E4 proteins as in (43). All viruses were purified by double CsCl banding and viral particle (vp) concentrations were calculated from OD260.

### Mice

Female C57BL/6 were purchased from Charles River Laboratories (Wilmington, Massachusetts, USA). C57BL/6-CD46 transgenic mice were generously provided by Dr. Roberto Cattaneo and Dr. Kah Whye Peng at Mayo Clinic. Mice were housed in the Mayo Clinic Animal Facility under the Association for Assessment and Accreditation of Laboratory Animal Care (AALAC) guidelines with animal use protocols approved by the Mayo Clinic Animal Use and Care Committee. All animal experiments were carried out according to the provisions of the Animal Welfare Act, PHS Animal Welfare Policy, the principles of the NIH Guide for the Care and Use of Laboratory Animals, and the policies and procedures of Mayo Clinic.

### Cryo-Electron Microscopy of Ad26

We recently described the first structure of species D Ad26 by cryo-electron microscopy (cryo-EM) at 3.7Å resolution and details of these analyses can be found in (26). Figure 1A shows the radially color-coded Ad26 virion down the icosahedral 2-fold axis and figure 1B shows a representation of the short-shafted Ad26 fiber and penton base using coordinates from (26).

### *In Vitro* Virus Infections

Cells were plated in 96-well black cell culture plates and infected at 50% confluence with the indicated viruses. Cells were infected at indicated multiplicities of infection (MOI) in terms of virus particles/cell. At indicated time points, media was removed and Bright-Glo luciferase reagent (Promega, Madison, WI) diluted 1:1 with PBS was added and luciferase activity was measured using the Beckman Coulter DTX 880 Multimode Detector system. The indicated viral particles were then added to the well for 48 hours before luciferase assay.

### Injections in Mice

Animals were housed under the Association for Assessment and Accreditation of Laboratory Animal Care (AALAC) guidelines in the Mayo Clinic Animal Facility. Experiments were performed under animal use protocols approved by the Mayo Clinic Animal Use and Care Committee. All experiments were performed under the provisions of the Animal Welfare Act, PHS Animal Welfare Policy and the principles of the NIH Guide for the Care and Use of Laboratory Animals. C57BL/6 or C57BL/6-CD46 transgenic mice were injected a single time by the indicated routes with the indicated amounts of Ad5 or Ad26 virus expressing GFP-Luc or with PBS. For tumor studies, 106 B16-CAR-CD46 cells were injected subcutaneously in Matrigel. Tumors were injected with RC-Ad5-GL or RC-Ad26-GL when the average size of tumors reached 200 mm3. The mice were imaged at the indicated timepoints on a Xenogen IVIS200 to measure *in vivo* luciferase activity.

### CBCs and Blood Chemistry

C57BL/6-CD46 transgenic mice were injected with 10^11^ vp of RC-Ad26. After 24 hours, blood was collected via submandibular vein puncture and collected in BD lithium heparin and K2 EDTA tubes. Complete blood cell counts (CBCs) and blood chemistry were analyzed on VetScan HM5 and Piccalo Express instruments, respectively. Controls were completed following the same procedure except no virus was administered prior. These analyses were performed by Mayo Pharmacology and Toxicology Core.

### Statistical Analyses

All analyses were performed using GraphPad Prism.

## RESULTS

### Ad26 Infection via CD46 on Cells

New data demonstrates that Ad26 interacts with CD46 via its hexon protein rather than by its fiber protein (47). Yet the question remains as to how the virus might use and balance interactions with CD46, integrins or sialic acid in natural and therapeutic settings. To examine this further, replication-defective Ad26 (RD-Ad26) expressing GFP-Luciferase was incubated with Chinese hamster ovary (CHO) cells that naturally lack both CAR and CD46. This infection was compared to CHO-CD46-C1 cells that express the C1 variant of human CD46 that includes only the single C STP domain (54) and to CHO cells expressing human CAR and human Junctional Adhesion Molecule 1 (JAM1) (**Fig. 3**). In the absence of any added human receptors, Ad26 infected CHO cells at significant levels (p < 0.0001 by one-way ANOVA). This significant infection could occur by interactions with the hamster integrin RGD loop or by sialic acid interactions. Adding CAR or JAM1 did not increase this transduction over unmodified CHO cells. In contrast, Ad26 infected CHO-CD46-C1 cells 6 to 7-fold higher than all of the other cell lines (p < 0.0001). These data suggest that Ad26 infection is facilitated specifically by CD46.

**Figure 3.**
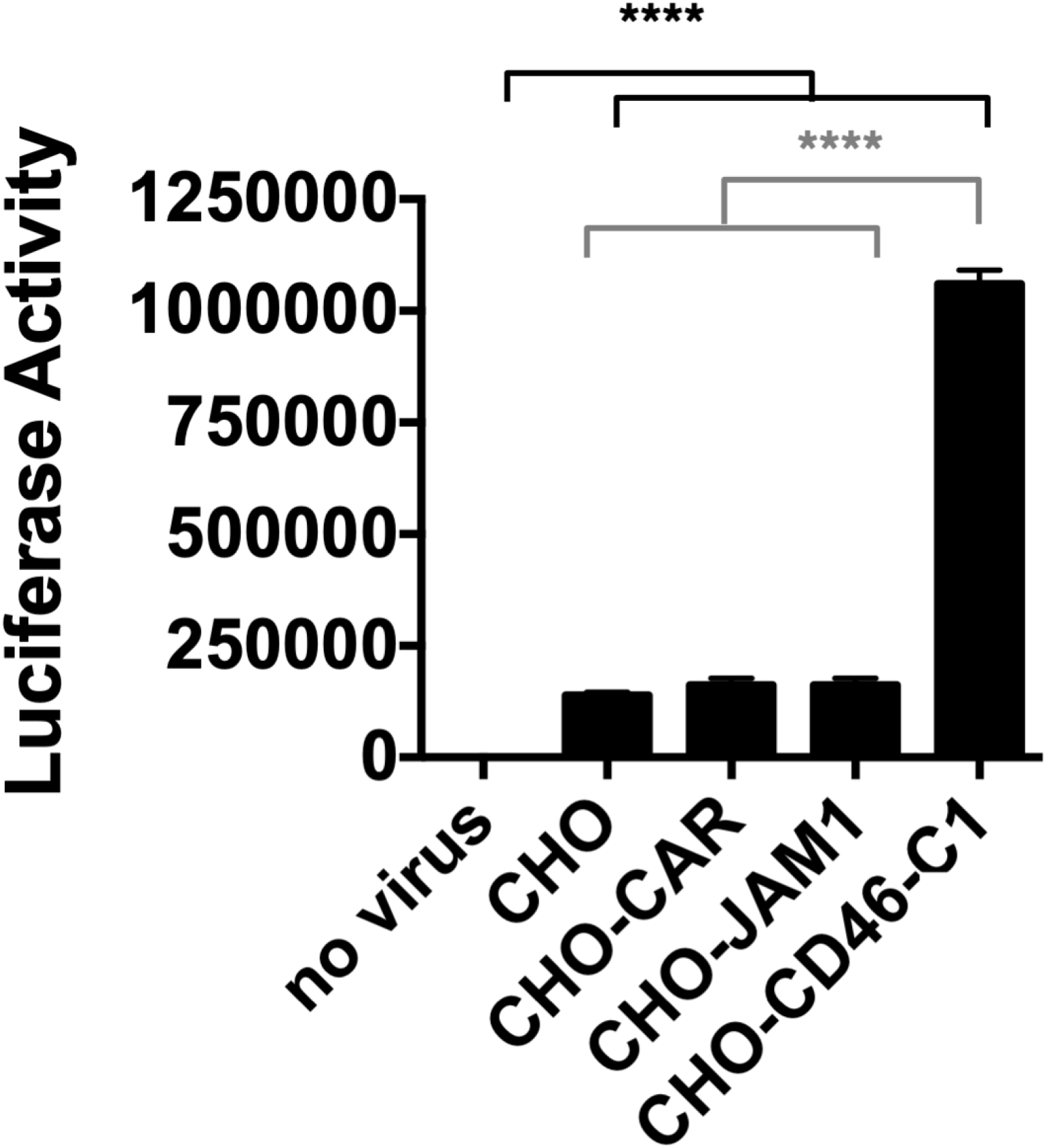
Infection of Ad26 on CHO Cells Displaying Different Human Receptors. RD-Ad26-GL expressing GFP-Luciferase was incubated with the indicated cells at a multiplicity of infection of 10,000 virus particles (vp)/cell and luciferase activity was measured 48 hours later. **** p < 0.0001 by one-way ANOVA.

### Ad26 Infection on Cells Expressing Truncated CD46 Variants

To test if, analogous to species B Ads, Ad26 functionally uses the membrane-distal CCP1 and CCP2 domains of CD46 (14, 19), Ad26 was applied to CHO cells expressing CD46-BC1 with deletions of CCP1, CCP2, or the membrane-proximal BC1-STP domain (55, 56). Ad26 infection was significantly lower on CD46-BC1-ΔCCP1 or ΔCCP2 (**Fig. 4**). Infection was also lower on CD46-BC1-ΔSTP cells (p < 0.001). These data suggest that Ad26 binds to CCPs 1 and 2 of CD46 and that the presence of an STP domain also influences this interaction.

**Figure 4.**
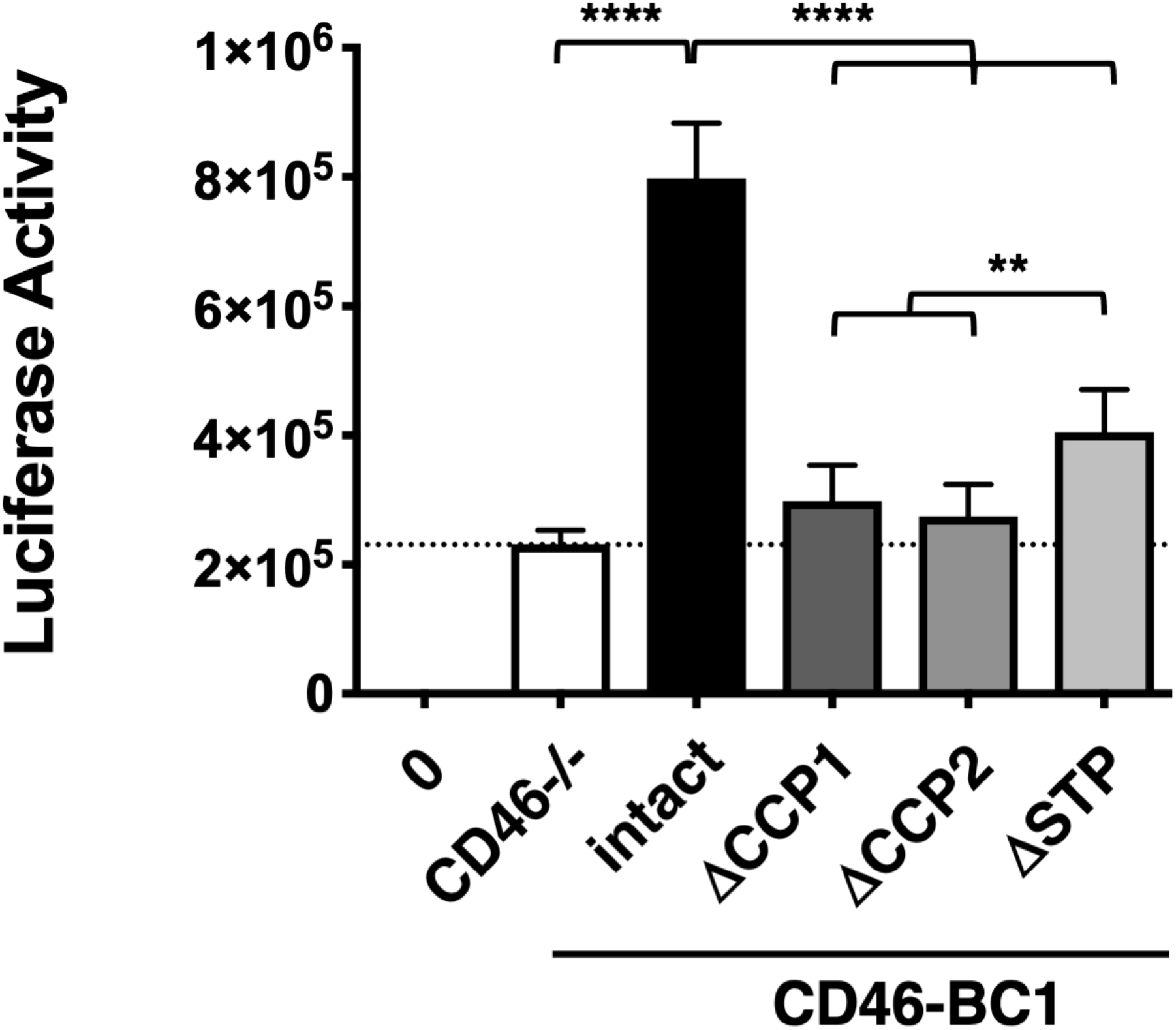
Infection of Ad26 on CHO Cells expressing CD46 with Deletions of Key Domains. ΔCCP1, ΔCCP2, and ΔSTP indicate CD46-BC1 proteins with deletions in these domains. CD46-R are negative control stable cell lines with CD46 in the antisense orientation to the expression cassette. Ad26-GL was incubated with the indicated cells at a multiplicity of infection of 10,000 virus particles (vp)/cell and luciferase activity was measured 48 hours later. ** p < 0.01, **** p < 0.0001 by one-way ANOVA.

### Ad26 Infection via CD46 with and without the Presence of Sialic Acid

We previously showed that a combination of CD46 antibody and cyclic RGD peptide can almost completely inhibit Ad26 infection of primary human B cells (7). In contrast, treating cells with neuraminidase or with wheat germ agglutinin to block sialic acid interactions did not inhibit Ad26 in our hands (7). This suggested that Ad26 uses CD46 and integrins, but not sialic acid, at least on these cells. In contrast, recent studies showed that Ad26 knob binds sialic acid at the same binding site as Ad37 (46).

Since the CCP1-2 and STP domains of CD46 are all heavily glycosylated, we hypothesized that Ad26 might not use CD46 as a protein receptor, but might instead use the protein as a scaffold that displays sialic acid (**Fig. 2** and (33)). In this model, over-expression of CD46 might simply increase the raw amount of sialic acid on the cell surface. Alternately, the extended “hockey stick” structure of CD46 (32) might simply provide an easily accessible landing pad to bind sialic acid on cells.

To test this, we expressed several versions of CD46-BC1 on either sialylation-competent CHO cells or sialylation-deficient LEC2 cells. LEC2 cells are CHO cell mutants that have a 90% reduction in sialylation of cellular glycoproteins due to an inability to translocate CMP-sialic acid across Golgi vesicle membranes, and thus result in different glycosylation patterns (**Fig. 5A**, right panel) (57, 58). In addition to wild type CD46-BC1, we generated constructs in which the complete ectodomain (Ecto) from CD46-BC1 (containing the 4 CCPs and 2 STP domains) were fused to a heterologous transmembrane domain from platelet-derived growth factor receptor (PDGFR). This allows display of all of the ecto domain of CD46, but fusion to PDGFR also removes the CD46 cytoplasmic domain that may also provide intracellular signaling functions (59). Another construct was made displaying only the -STP domain from BC1 fused to PDGFR (STP) to display only this heavily glycosylated domain without native CD46 intracellular domain.

**Figure 5.**
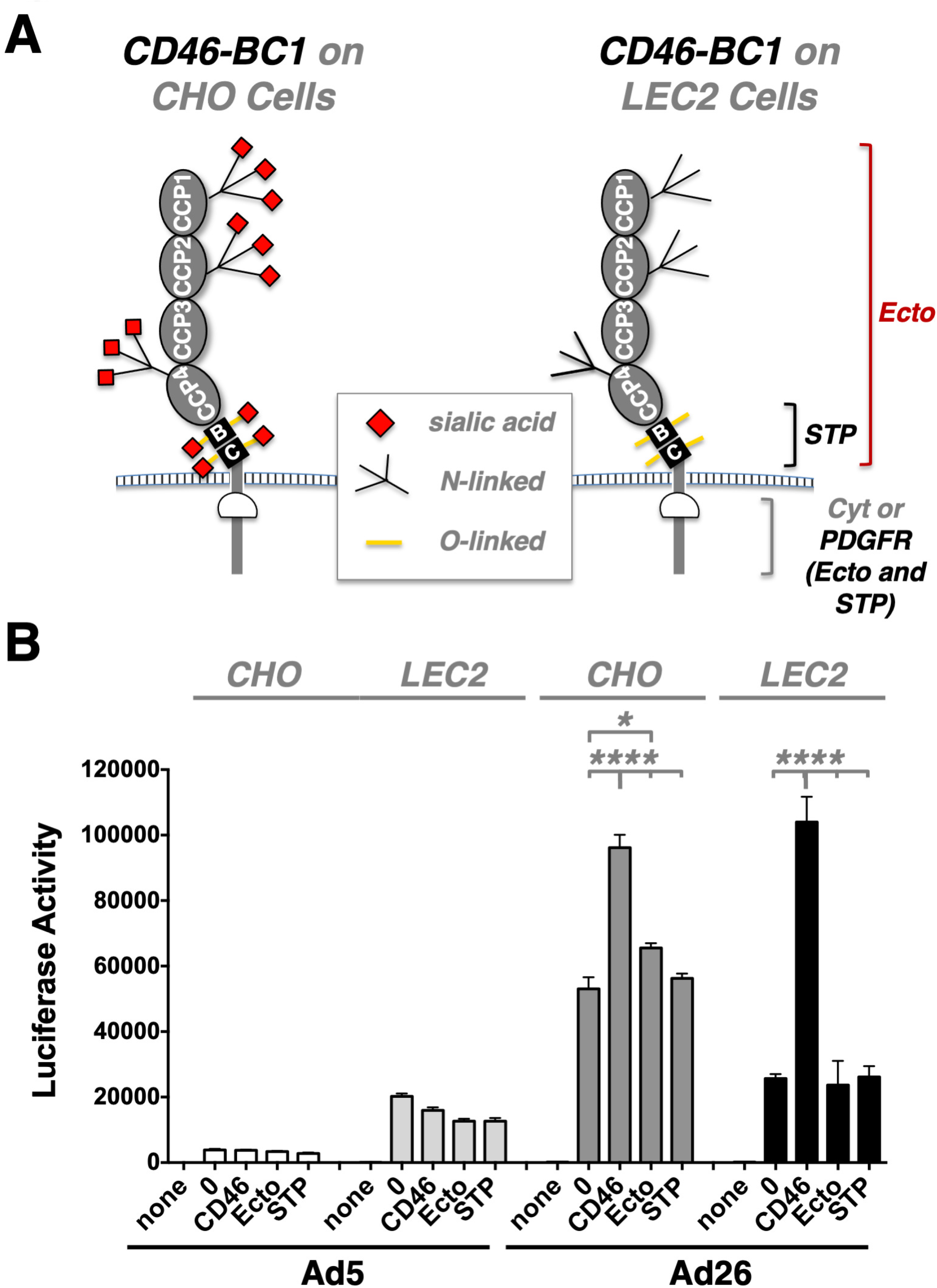
Effects of Sialylation on Ad26 Infection of CD46 Mutant Cells. **A)** Schematic of CD46 glycosylation in CHO and LEC2 sialylation-deficient cells. **B)** The indicated cells were incubated with Ad26-GL and luciferase activity was measured 2 days later. * p < 0.05, **** p < 0.0001.

When Ad5 and Ad26 were used to infect CHO cells expressing these proteins, Ad26 overall mediated significantly stronger gene delivery than CAR-utilizing Ad5, irrespective of whether CD46 was present or not (**Fig. 5B**). However, Ad26 once again infected CHO cells expressing wild type CD46-BC1 cells more efficiently than unmodified CHO cells (p < 0.0001). Interestingly, Ad26 infection of the STP-PDGF and Ecto-PDGF chimeras was no better than the transduction of unmodified CHO cells.

The same CD46 plasmids were used to modify LEC2 cells (57). Indeed, Ad26 transduction of unmodified LEC2 cells resulted in more than 50% decreased luciferase levels compared to CHO cells – as was the case for LEC2 cells expressing Ecto-PDGF and STP-PDGF, which again yielded no discernable effect. However, when LEC2 cells overexpressing wild type CD46-BC1 were infected with Ad26, the level of luciferase gene expression was virtually identical to that observed on CHO-CD46-BC1 cells (**Fig. 5B**). These results suggested that Ad26 does use sialic acid for some level of infection, perhaps on other cellular proteins, but that CD46 can functionally substitute sialic acid when expressed at high levels, irrespective of its sialylation status. Ad26 clearly does not use CD46 as a mere scaffold for carbohydrate binding, and surprisingly, its transmembrane and cytoplasmic domains seem to be relevant for Ad26 transduction as well. Ad5 infection interestingly increased on all of the LEC2 cells.

### *In Vivo* Infection by Ad26 in Normal and CD46 Transgenic Mice

CD46 is expressed on nearly all human cells (28). In contrast, normal mice express CD46 only in the testes and retina (39-41). We previously showed that *in vivo* intramuscular (IM) injection of RD-Ad26 into C57BL/6-CD46 transgenic mice mediated significantly stronger transduction than similar injections in normal BALB/c mice (43). While this supports the use of CD46 by the virus *in vivo*, this was not a direct comparison of Ad26 in mice with and without CD46.

To examine this more directly with viruses that are relevant to oncolytic virotherapy, replication-competent Ad5 (RC-Ad5) and replication-competent Ad26 (RC-Ad26) expressing GFP-Luciferase were compared *in vivo* in C57BL/6 or C57BL/6-CD46 transgenic mice (60) (**Figure 6**). These C57BL/6-CD46 transgenic mice contain a knock-in of the human CD46 gene as well as locus control regions from human chromosome 1, granting these mice near-ubiquitous expression of the human CD46 protein with expression patterns dictated by the human promoter sequence (56). To control for heterogeneity of mouse genetic backgrounds skewing transduction, the C57BL/6 mice used for these experiments were CD46-littermates of the CD46^+^ mice used.

**Figure 6.**
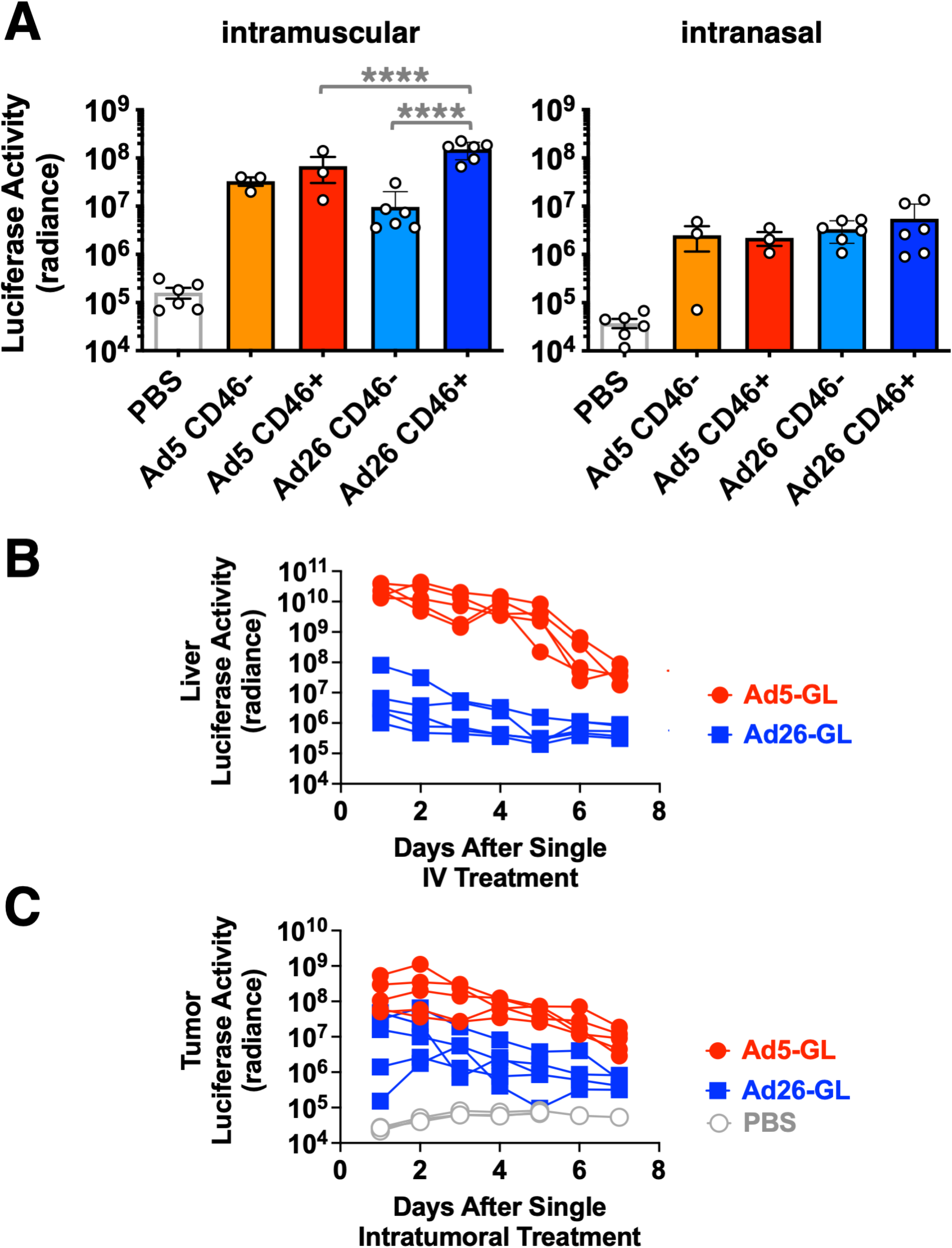
*In Vivo* Luciferase Activity After Intramuscular, Intranasal, Intravenous, and Intratumoral Administration of Ad5 or Ad26 in CD46^-^ and CD46^+^ Mice. **A)** Luciferase imaging 2 days after intranasal or intramuscular injection of 1×10^10^ vp of RC-Ad5-GL or RC-Ad26-GL. **B)** Luciferase activity in livers 2 days after intravenous injection of 3×10^10^ vp of RC-Ad5-GL or RC-Ad26-GL into CD46 transgenic mice **C)** Luciferase activity in tumors 2 days after intratumoral injection of 3×10^10^ vp of RC-Ad5-GL or RC-Ad26-GL into B16-CAR-CD46 subcutaneous tumors. Graphs show the same data on a log scale on the left and a linear scale on the right. * p < 0.05, **** p < 0.0001.

1×10^10^ virus particles (vp) of RC-Ad5-GL or RC-Ad26-GL were injected intramuscularly (IM), intranasally (IN), intravenously (IV), or intratumorally (IT) into groups of 5 mice and luciferase imaging was performed 2 days later (**Fig. 6**). Ad26 mediated 15-fold higher transduction in CD46^+^ mice than CD46-mice by the IM route (p < 0.0001 by one-way ANOVA, **Fig. 6A**). Ad26 transduction was also 2-fold higher than Ad5 transduction in the muscle of CD46^+^ mice. In contrast, Ad5 and Ad26 transduction was nearly equal in mice with and without CD46 when delivered intranasally. Overall, transduction by both vectors decreased approximately 10-fold by the IN route when compared to the IM route. We therefore conclude that the usage of CD46 by Ad26 *in vivo* is dependent on the administered route, and that there seems to be preferred usage of CD46 following IM administration.

RC-Ad5-GL and RC-Ad26-GL were next injected into B16-CAR-CD46 subcutaneous tumors in CD46 transgenic mice and luciferase expression was measured (**Fig. 6B and C**). After intravenous administration, Ad5 mediated more than 100-fold higher luciferase expression in the liver than Ad26 (**Fig. 6B**). When the two vectors were injected IT into the B16-CAR-CD46 tumors, Ad5 mediated up to 10-fold higher tumor infection than Ad26 (**Fig. 6C**). While this gives no indication on the specific usage of CD46, the results show how two different vectors can have vastly different efficacies depending on how they are administered.

### Complete Blood Counts and Blood Chemistry after Intravenous Injection of Ad26 into CD46 Transgenic Mice

A low frequency of thrombosis and thrombocytopenia have been reported after COVID-19 vaccination with the ChAdOx1 nCoV-19 vaccine (Oxford–AstraZeneca) and the Ad26.COV2.S vaccine (Johnson & Johnson/Janssen) (48, 49). While the COVID-19 vaccines have been injected intramuscularly, some fraction of injected vaccine will leak into the blood and could potentially induce effects on the clotting system.

We previously examined in detail the ability of Ad5 to provoke clotting side effects after intravenous injection (61). An IV dose of 10^11^ vp of Ad5 induces significant thrombocytopenia within 24 hours. This response is rapid as demonstrated by increases in d-dimer fibrin degradation products in the blood within 6 hours of injection. Ad5 appears to induce this effect by binding and activating platelets and endothelial cells directly (61).

To examine if leak of Ad26 into the blood might also perturb the clotting system, a similar large 10^11^ vp dose of RC-Ad26 was administered IV to CD46 transgenic mice and complete blood counts (CBC) and blood chemistry analyses were performed one day later (**Fig. 7**). In contrast to Ad5, the IV bolus of Ad26 failed to provoke notable changes in platelets when compared to control animals (**Fig. 8A**).

**Figure 7.**
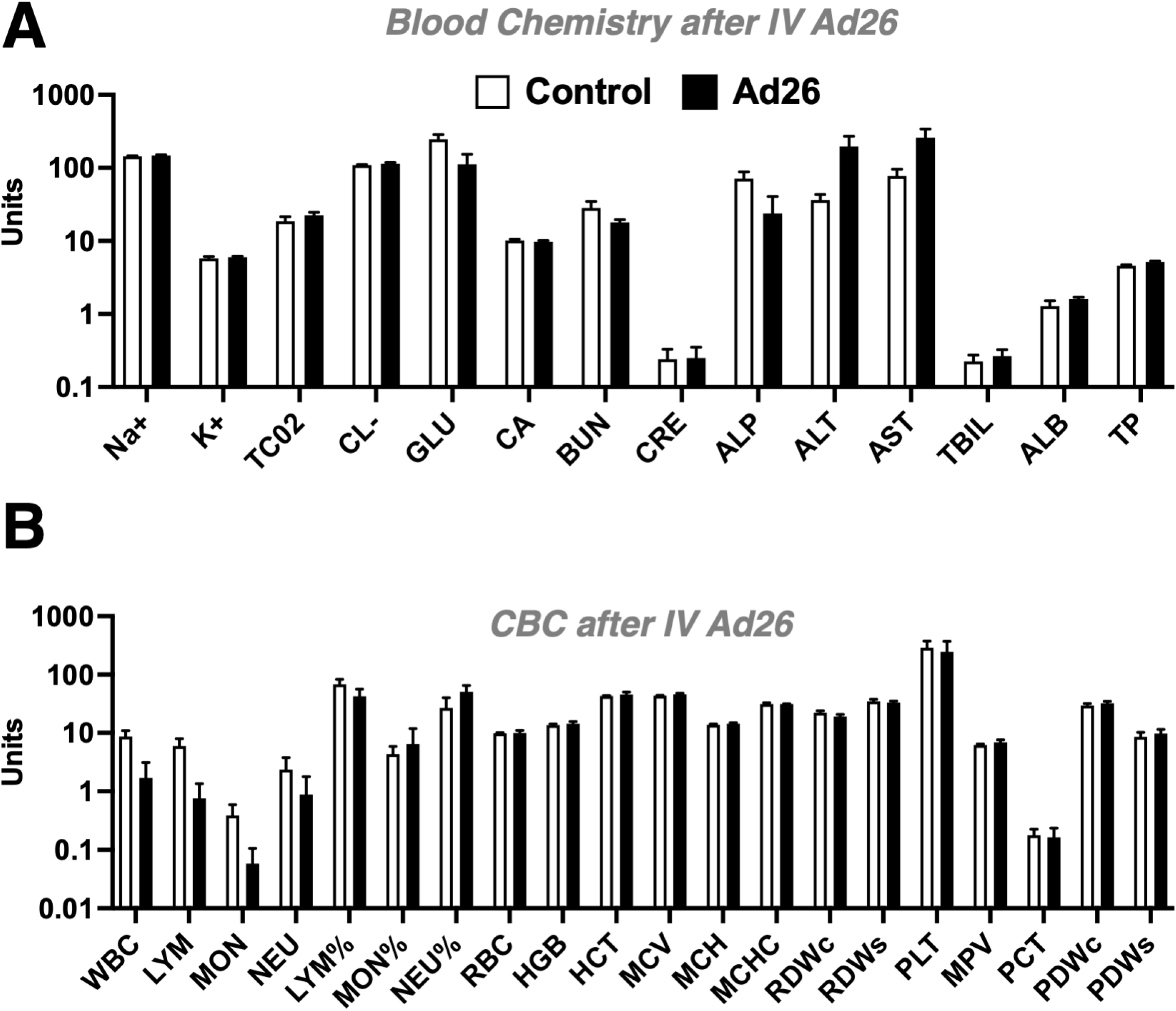
Effects of Intravenous Administration of Ad26 in CD46 Transgenic Mice on CBCs and Blood Chemistry. Groups of 5 CD46 transgenic mice were injected with a high dose of 1×10^11^ vp of RC-Ad26-GL and blood was collected one day later for CBC and blood chemistry analysis. Control values were collected from 5 uninjected CD46 mice.

**Figure 8.**
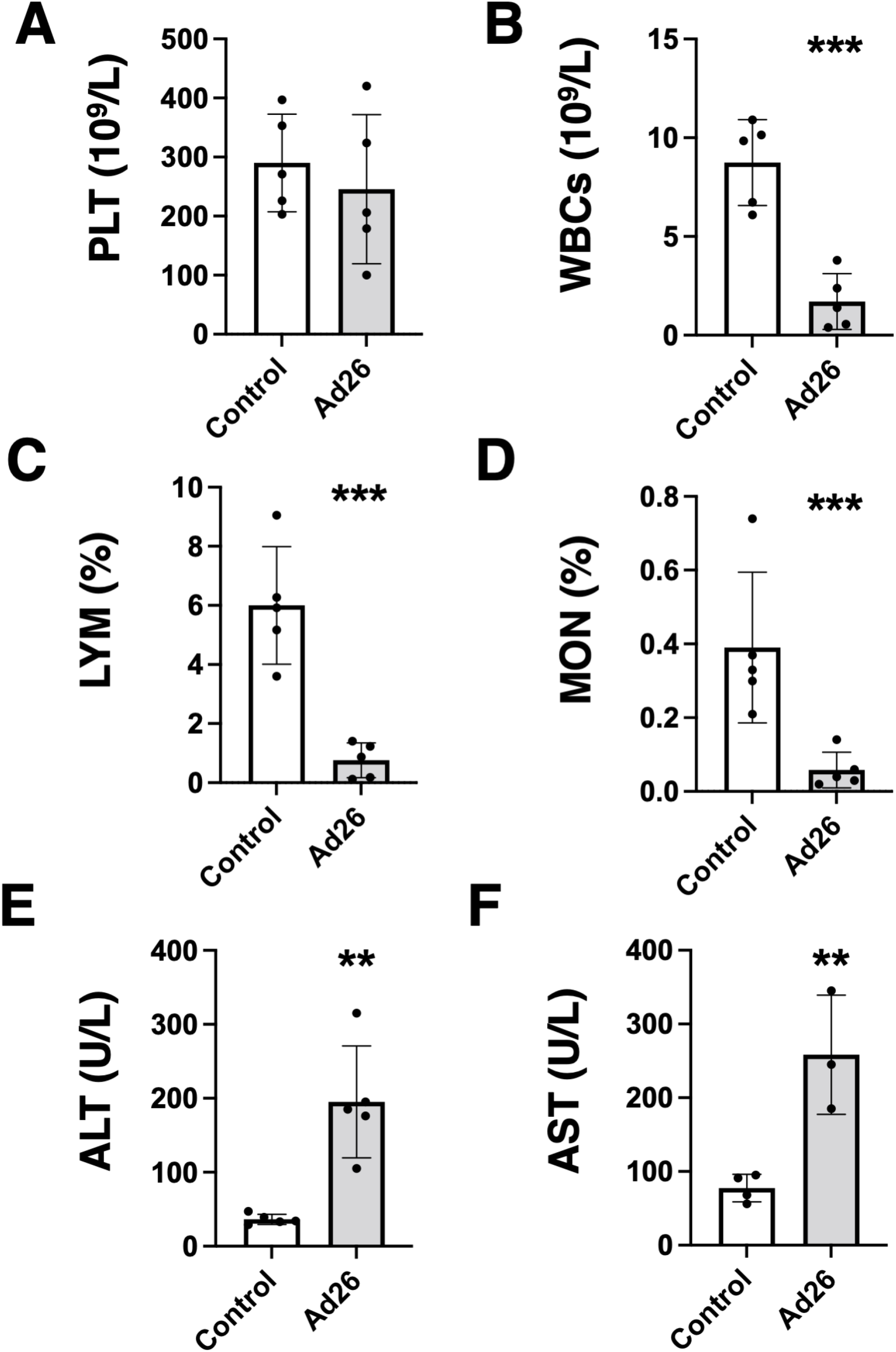
Effects of Intravenous Administration of Ad26 in CD46 Transgenic Mice on Platelets, White Blood Cells, and Liver Enzymes. Selected parameters from Figure 7 are shown for **A)** Platelets (PLT), **B)** White Blood Cells (WBCs), **C)** Lymphocytes (LYM), **D)** Monocytes (MON), **E)** Alanine aminotransferase (ALT), **F)** Aspartate aminotransferase (AST). ** p < 0.01, *** p < 0.001 by T test.

In contrast, Ad26 did induce significant reductions in white blood cells, lymphocytes, and monocytes (p < 0.001, **Fig. 8B, C, and D**) after IV injection. Ad26 also produced significant elevations in liver enzymes including alanine transaminase (ALT) and aspartate aminotransferase (AST) after IV injection (p < 0.01, **Fig. 8E and F**). (62)These data suggest that direct exposure of blood proteins and cells does not provoke thrombocytopenia but does cause reductions in certain nucleated white blood cells and liver damage.

## DISCUSSION

Our previous work found that species D human Ads can be potent oncolytic viruses against B cell cancers (6, 7) and have utility as mucosal gene-based vaccines (9, 43). Receptor utilization testing on primary patient B cell cancer cells showed that Ad26 uses CD46 and integrin binding to infect these cells (7). Blocking CD46 antibodies reduced Ad26 infection in these B cells, but this inhibition was not 100%. Combined blockade of CD46 and integrin could inhibit infection nearly 100% (7). Removing or blocking sialic acid on B cells had no effect.

More recent work reported that Ad26 knob does not bind CD46 and instead uses αvβ3 integrin as its primary receptor on epithelial cells (44). Other work showed that Ad26 knob could bind CAR, but could not bind CD46 (45).

Given these conflicting results and analyses on different cell types, this study was performed to examine the functional use of receptors by Ad26 in different circumstances. During the course of our work, another group published seminal work showing as we did that Ad26 does indeed use CD46, but not with its fiber, but rather with its hexon (47).

We proceeded to functional testing on more controlled target cells than the primary B cell cancer cells used in our previous studies, and to permutate CD46 constructs that lack specific parts of the protein. Ad26 testing on well-defined CHO-CD46 cell lines (28, 29, 33) showed that Ad26 infected cells that expressed intact forms of CD46. Ad26 infected cells expressing CD46 lacking CCP1 and CCP2 significantly less efficiently than wild-type CD46. Ad26 infection was also less efficient when CD46 lacked its STP or transmembrane and cytoplasmic domain. Thus, like some species of adenovirus types B and D, Ad26 binds CD46 via CCPs 1 and 2, yet is also influenced by the STP and transmembrane/cytoplasmic domains. It is currently unclear whether this influence involves direct or allosteric effects.

However, CD46 (over)expression on CHO cells only increased Ad26 infection 6-fold. This weak utilization of CD46 is consistent with earlier observations that Ad26 infects CD46-expressing cells only half as well as known CD46-targeting Ad11 and Ad35 (3).

When we tested Ad26 *in vivo* in CD46^-^ and CD46^+^ transgenic mice, Ad26 infected CD46^+^ mice 15 times higher than CD46^-^ littermates after intramuscular injection. However, CD46 expression had little effect on Ad26 by the intranasal route. This suggests that Ad26 interactions with CD46 may be important during intramuscular vaccination but may be less relevant during natural respiratory infections.

When Ad5 and Ad26 were put to the stringent test of transduction after intravenous injection, Ad26 was 100 to 100,000-fold less effective than Ad5 at transducing the liver or other sites. When the two vectors were injected intratumorally into CD46-expressing B16 melanoma tumors to model oncolytic therapy, Ad5 was again more robust than Ad26.

Considering observations of clotting disorders during Ad26 COVID-19 vaccinations, we examined whether RC-Ad26 might provoke similar effects in CD46 transgenic mice in a worst-case scenario of injecting the virus directly into the bloodstream. Ad26 failed to provoke notable changes in platelets, but did induce significant reductions in white blood cells, lymphocytes, and monocytes. This is consistent with observations that RC-Ad26 can reduce the viability of human CD14+ monocytes and CD3+ lymphocytes *ex vivo* (7). Ad26 also produced significant liver damage in the CD46 transgenic mice as demonstrated by elevating liver enzymes without effectively transducing the liver. These data suggest that Ad26 virions themselves do not appear to directly induce thrombocytopenia, but that it does damage white blood cells and hepatocytes at least when delivered at high doses as a replication-competent virus.

In summary, these data indicate that Ad26 can functionally interact with CD46 for infection *in vitro* and *in vivo*. It should be noted that these are observed under conditions where CD46 is ectopically expressed in cells or in mice. While this is ectopic expression in mice, this expression is driven by the natural locus control regions for CD46 from human chromosome 1, so should mimic to some degree expression in human tissues.

While it is clear the Ad26 can use CD46 *in vivo*, this appeared important only by the intramuscular route. This is important for the virus’ efficacy as an intramuscular vaccine, but less relevant during natural respiratory mucosal infections. Understanding the *in vivo* tropism of Ad26 is important not only for considering vaccine efficacy, but also vaccine safety. Further examination of Ad26’s ability to interact with CD46 and other receptors is warranted to better understand the implications of these interactions for its effective use in vaccines and therapies.

## ACKNOWLEDGEMENTS

This project was supported by a Developmental Project to M.A.B. from the University of Iowa/Mayo Clinic Lymphoma Specialized Program of Research Excellence (SPORE) in Lymphoma (P50 CA097274) and the Walter & Lucille Rubin Fund in Infectious Diseases Honoring Michael Camilleri, M.D. at Mayo Clinic.

